# Convergent losses of arbuscular mycorrhizal symbiosis in carnivorous plants

**DOI:** 10.1101/2025.04.03.646726

**Authors:** Héctor Montero, Matthias Freund, Kenji Fukushima

## Abstract

Across evolutionary scales, lineages acquire and lose traits and associated genes. Most land plants form arbuscular mycorrhizal (AM) symbiosis, an ancient trait for enhanced nutrition that was convergently lost in some clades. Carnivory, another nutritional trait, is a more recent adaptation that has convergently arisen in several angiosperm orders. The two biotic interactions similarly help plants acquire mineral nutrients, raising the question of whether they can coexist. However, the mycorrhizal status of carnivorous plants has long remained speculative. Here, we surveyed the occurrence of AM-associated genes in five angiosperm orders harbouring carnivorous species, revealing convergent losses of the AM trait either coincident with or predating the emergence of carnivory. Exceptionally, the carnivorous plant species *Roridula gorgonias* retains symbiosis-related genes and forms arbuscules upon inoculation assays, demonstrating the two nutritional strategies, although rare, can coexist. The youngest carnivorous lineage, *Brocchinia reducta*, showed signatures of the early stages of AM trait loss, as reflected by its gene retention and AM colonization patterns. An AM-associated *CHITINASE* gene encodes a digestive enzyme in the Australian pitcher plant *Cephalotus*, suggesting gene co-option. These findings illuminate the largely unexplored processes by which plant nutritional strategies evolve and supplant one another over time.

## Introduction

Nutrient acquisition is pivotal for plant fitness, and evolution is expected to select for traits that efficiently modulate this process. Plants are able to capture mineral nutrients directly from their environment and by interacting with other organisms.

Most land plants form a mutualistic relationship known as the arbuscular mycorrhizal (AM) symbiosis with fungi from the subphylum Glomeromycotina. This is an ancient interaction that is believed to have enabled plant transitions from aquatic to terrestrial habitats and that accompanied early land plants, predating in millions of years the evolution of roots or flowers (Strullu-Derrien et al. 2018; Rich et al. 2021). In return for photosynthesis-fixed carbon, AM fungi provide plants with mineral nutrients such as phosphorus, nitrogen and potassium (Lanfranco et al. 2018).

During the course of land plant evolution, several lineages have independently lost the ability to establish symbioses with AM fungi. This scenario is usually accompanied by plants partnering with other root symbionts or by evolving alternative nutrient acquisition strategies including parasitism and the formation of specialized roots adapted for phosphate scavenging such as those in the families Proteaceae and Cyperaceae. The non-AM trait is also found among plants that lack developed underground roots including some epiphytes and aquatic species (Lambers and Teste 2013). Cases also exist where AM symbiosis is absent in plants with seemingly ordinary lifestyles, notably in the Brassicaceae (Sharma et al. 2023). Phylogenomic approaches have been used to identify AM symbiosis-associated genes (hereafter symbiosis genes) that are lost upon mutualism breakdown (Delaux et al. 2014; Favre et al. 2014; Bravo et al. 2016). Exploring the evolutionary paths leading to the loss of the AM symbiosis would be beneficial for better understanding how plants optimize their nutrient acquisition strategies. Plant carnivory is a convergently evolved trait that is evolutionarily younger than AM symbiosis and has arisen in six angiosperm orders (Fleischmann et al. 2018; Lin et al. 2021). Carnivory takes place mainly by means of highly specialized leaves that form traps, capable of capturing, killing, and digesting arthropod prey. The nutrients acquired from prey are of similar nature than those that plants acquire from AM fungi. This posits questions on functional redundancy between the two nutritional traits, AM symbiosis and carnivory. However, research on biotic interactions in carnivorous plants usually centers more on prey than on root biology (Montero and Fukushima 2023), and whether or not carnivorous plants can form mycorrhizae is uncertain. Past studies (see below) aimed to analyse their root microbiota in natural communities with equivocal outcomes, and no study has attempted inoculation assays. One study reported the presence of AM fungal DNAs in an attempt to analyze the root endophytes in the flypaper-type carnivorous plant *Drosera rotundifolia* (Quilliam and Jones 2010). Another study made microscopic examinations in this species and found aseptate hyphae and vesicles, but no arbuscules (Weishampel and Bedford 2006). Reports also exist on low colonization and presence of arbuscules in *D. intermedia*, but no microscopy image is provided (Fuchs and Haselwandter 2004). In another publication, arbuscules are said to occur in *D. burmannii* and *D. indica* (Harikumar 2013), but the images provided do not show arbuscules and may not correspond to AM fungi. Coincidently, all these studies had made *Drosera* their study subject, a group of carnivorous plants in the Caryophyllales. Dedicated studies in other carnivorous plants are lacking. Recent research compiles these incomplete observations into a large-scale dataset to derive evolutionary insights (Werner et al. 2018). Altogether, the mycorrhizal status of carnivorous lineages lacks unequivocal evidence, with the unsubstantiated notion of them being non-mycorrhizal (Adlassnig et al. 2005).

The convergent evolution of carnivory provides an excellent opportunity to test the patterns of AM symbiosis loss associated with the emergence of alternative nutritional strategies. In order to scrutinize possible connections between the independent losses of AM symbiosis and the independent gains of plant carnivory, we undertook a comparative genomics approach to explore the presence of symbiosis-related genes in carnivorous plants across angiosperms. Additionally, we coupled this with inoculation assays and microscopic examinations. Our analyses suggest that AM symbiosis has been independently lost in most, but not all, carnivorous lineages.

## Results

### Occurrence of symbiosis genes in carnivorous plants

In order to determine the occurrence of symbiosis-related genes in carnivorous lineages, we generated a dataset of 124 genomes and 105 transcriptomes centred on five plant orders where carnivory evolved and representatives in the rest of angiosperm orders. Publicly available data was supplemented by a set of newly generated transcriptomes from multiple tissues including roots. We identified the orthologs of the 75 symbiosis genes, which were previously identified through a phylogenomic method (Bravo et al. 2016). First, we employed two distinct approaches for orthogroup assignment: a graph-based tool, SonicParanoid2 (Cosentino et al. 2024), and a BLAST sequence search combined with phylogenetic orthogroup identification (see Methods). Next, we reconstructed a species-tree-aware maximum-likelihood gene tree for each orthogroup to accurately detect orthologous relationships, even in cases of complex evolutionary histories involving lineage-specific gene duplications and losses. Finally, we reported the occurrence of symbiosis genes if one or more orthologs were detected by either or both orthogroup construction methods. Genome sequencing, gene prediction, and transcriptome assembly do not guarantee the complete recovery of gene sets, making it difficult to distinguish between genuine gene loss and technical omissions for individual genes. However, gene set completeness scores were high for genomes (BUSCO completeness = 89.5±8.6%, mean ± standard deviation, *N* = 124 species) and transcriptomes (77.1±18.1%, *N* = 105), indicating that our dataset provides sufficient gene coverage to capture gene retention trends across multiple species and genes.

Altogether, we compiled data from species belonging to 46 out of the 64 recognized angiosperm orders (The Angiosperm Phylogeny Group et al. 2016), a substantial increase compared to previous phylogenomic studies on AM symbiosis, which reached 18 to 20 angiosperm orders (Bravo et al., 2016, Radhakrishnan et al., 2021). Focusing on the orders where carnivory evolved, we found that most symbiosis genes were undetected in the majority of carnivorous plants, suggesting convergent gene losses (Fig. 1 and Supplementary Fig. 1). In the Lamiales, symbiosis genes were mostly undetected in all four examined carnivorous genera in the Lentibulariaceae and the Byblidaceae while they tended to be preserved in the other lineages, including the most closely related non-carnivorous species in the dataset. A similar situation occurred in the Oxalidales; the carnivorous *Cephalotus follicularis* (Cephalotaceae) lacked 80% (60/75) of the symbiosis genes while starfruit (*Averrhoa carambola*, Oxalidaceae), the phylogenetically closest genome in the dataset, had a nearly complete repertoire. As such, it appears that the emergence of carnivory coincides with the loss of AM symbiosis in *C. follicularis* albeit it is difficult to assess which trait change occurred first as this is the only species in this carnivorous lineage. In the Caryophyllales, all five carnivorous plant genomes lacked the symbiosis genes. In this case, however, we found all the species in the order in our dataset to be devoid of the symbiosis genes. In the Ericales, symbiosis genes were undetected in the transcriptomes of the carnivorous species in the Sarraceniaceae belonging to the genera *Darlingtonia*, *Heliamphora* and *Sarracenia*. Although transcriptomes generally exhibit lower gene coverage reliability compared to genomes, the symbiosis genes tended to be consistently either detected or undetected across the three genera, suggesting robust gene detection. The symbiosis genes retained in all examined Sarraceniaceae species include, for example, the *SYNTAXIN OF PLANTS132* (*SYP132*) (Huisman et al. 2016; Pan et al. 2016; Liu et al. 2022) and the cuticular wax-related gene *DROUGHT-INDUCED WAX ACCUMULATION 1* (*DWA1*) (Zhu and Xiong 2013). In contrast, the genome of the flypaper-type carnivorous plant *Roridula gorgonias*, from the independently evolved carnivorous family Roridulaceae, surprisingly presented most of the symbiosis genes (67/75). Finally, in the Poales, we found another exception where symbiosis genes were well detected in a carnivorous plant, corresponding to the transcriptome of the pitfall-type carnivorous plant *Brocchinia reducta* (Bromeliaceae). Although transcriptomes are usually less complete than high-quality genome assemblies in terms of gene set coverage, we found 67% (50/75) of symbiosis genes in this species. These include canonical components of the common symbiosis signalling pathway, such as *SYMBIOSIS RECEPTOR-LIKE KINASE* (*SYMRK*), *CASTOR*, *CALCIUM AND CALMODULIN-DEPENDENT PROTEIN KINASE* (*CCaMK*) and *CYCLOPS* (Parniske 2008), and also signalling-related genes known to function in arbuscule-containing cells, such as *ARBUSCULE DEVELOPMENT KINASE 1* (*ADK*) (Guo et al. 2022; Shi et al. 2022), *ARBUSCULAR RECEPTOR-LIKE KINASE 1* (*ARK1*) (Roth et al. 2018; Irving et al. 2022), *CYCLIN-DEPENDENT KINASE-LIKE (CKL*) (Ivanov and Harrison 2024) and *RECEPTOR-LIKE CYTOPLASMIC KINASE 171* (*RLCK171*) (Leng et al. 2023).

**Figure 1.**
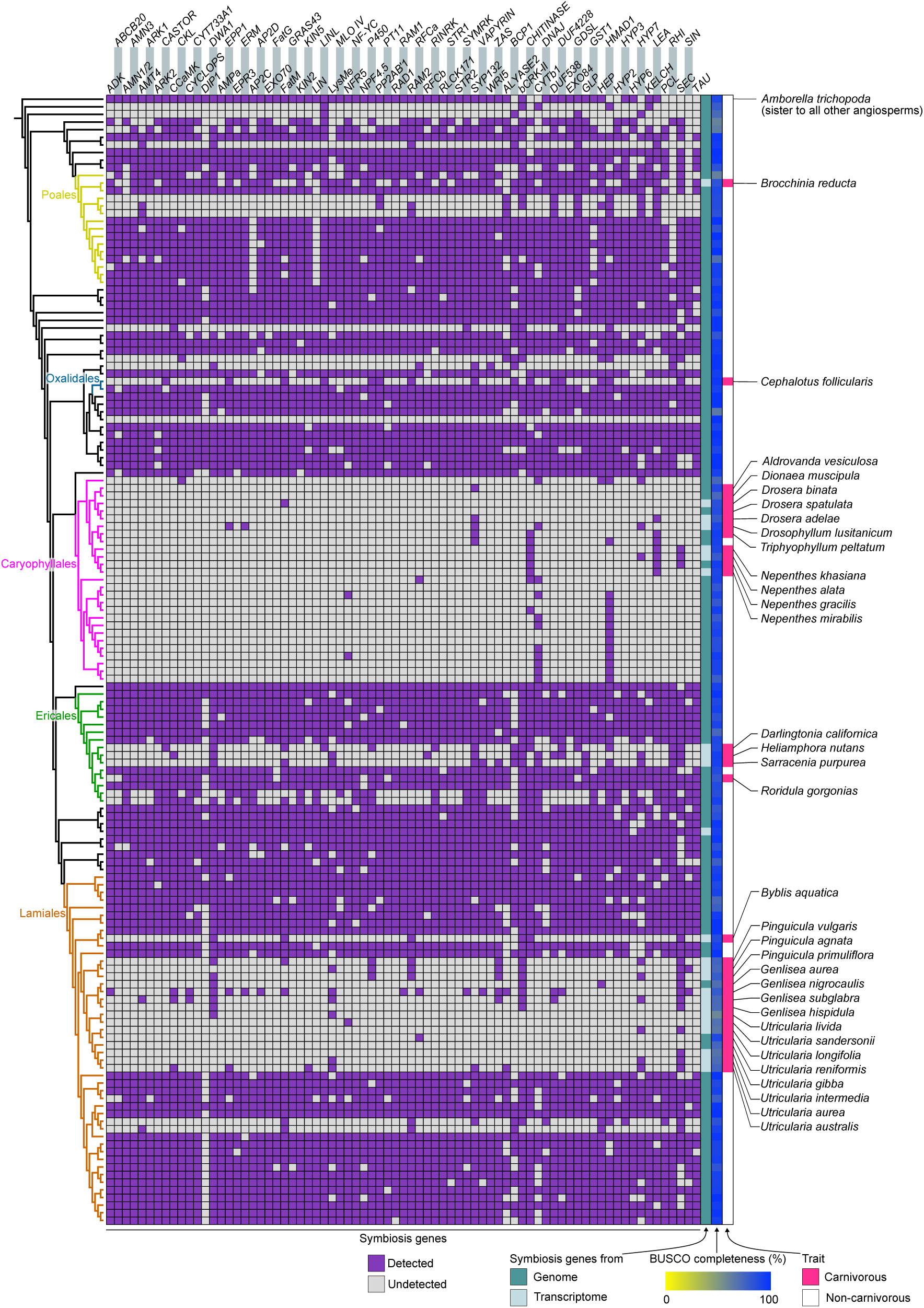
Occurrence of symbiosis genes in carnivorous plants. Symbiosis genes present across angiosperms. Occurrence reflects the consensus between orthogroups and Blast-based analyses. Transcriptomes of non-carnivorous plant species are excluded here. For the complete dataset, see Supplementary Figure 1.

Besides carnivorous plants, there were certain plant lineages where symbiosis genes were not detected. In the Poales, sedges and rushes from the genera *Juncus*, *Rhynchospora* and *Carex*, lacked most symbiosis genes but also commonly retained some such as *DOMAIN OF UNKNOWN FUNCTION 538* (*DUF538*). In the Ericales, species in the family Ericaceae, known to engage in an alternative mycorrhizal association called Ericoid mycorrhizal symbiosis (Smith and Read 2008), had retained around 45% of the symbiosis genes in autotrophic species (42–53%, *N* = 3 species). The transcriptomes of two mycoheterotrophic species in this family, *Monotropa hypopitys* and *Monotropastrum humile*, presented 18 and 21% of the symbiosis genes, respectively. In the Lamiales, parasitic species in the Orobanchaceae lacked most symbiosis genes, albeit retaining a few such as the known AM lipid biosynthesis enzymes *FatM* and *REQUIRED FOR ARBUSCULAR MYCORRHIZA 2* (*RAM2*) (Fig. 1).

To explore the contribution of AM symbiosis loss in the overall convergent gene loss in carnivorous plants, we expanded our focus from targeted analyses of known symbiosis genes to the entire orthogroup dataset, asking which genes are commonly lost in carnivorous plant genomes. To this end, we identified orthogroups that are absent from all the carnivorous plant genomes and transcriptomes in the Oxalidales and Lamiales, yet present in representative non-carnivorous genomes of those orders. We selected these two orders from the five analyzed orders for several reasons. Firstly, there are no available carnivorous genomes in the Poales. Secondly, the Caryophyllales atypically lacks symbiosis genes in both carnivorous and non-carnivorous lineages. Thirdly, the Ericales includes *Roridula gorgonias*, which maintains the symbiosis genes despite carnivory. Lastly, Oxalidales and Lamiales represent independent evolutionary origins of carnivory in the rosids and the asterids, the two major eudicot clades. Out of a total of 183,957 orthogroups, 11,274 were commonly present in the target non-carnivorous species, while 65,520 were commonly absent in the target carnivorous species. From these, 85 orthogroups fulfilled the criteria of being present in the non-carnivorous species and absent in the carnivorous species. Approximately one-third of these 85 orthogroups were associated with AM symbiosis, including 24 focal symbiosis orthogroups analysed in this study, as well as the AM symbiosis-related genes *ADK2a* (a paralogue of *ADK1*) (Guo et al. 2022), *AMMONIUM TRANSPORTER 2;3* (*AMT2;3*) (Breuillin-Sessoms et al. 2015), *MYB1* (Floss et al. 2017) and *MYCORRHIZA-INDUCED GRAS 1* (*MIG1*) (Heck et al. 2016) (Supplementary Fig. 2 and Supplementary Table 1). This result suggests that the loss of symbiosis genes is an important contributor to the convergent and potentially deterministic gene loss in carnivorous plants.

In summary, our dataset lacked the majority of symbiosis genes in 14 carnivorous genera, with the other two genera (i.e., *Brocchinia* and *Roridula*) being exceptions preserving most symbiosis genes. These results suggest that the evolution of plant carnivory tends to accompany the decay of molecular building blocks for AM symbiosis.

### AM inoculation assays in carnivorous plants

Given the presence of symbiosis genes in *R. gorgonias* and *B. reducta*, we performed AM inoculation assays to assess the ability of the plants to be colonized by AM fungi and develop arbuscules, the signature symbiotic structures diagnostic for nutrient exchange (Luginbuehl and Oldroyd 2017) (Fig. 2). Plantlets of *R. gorgonias* were inoculated with the AM fungus *Funneliformis mosseae*. Despite the difficulty in clearing *R. gorgonias* roots for fungal staining procedures, we successfully observed well colonized root zones with arbuscules at six weeks post-inoculation (wpi) (Fig. 2b). This shows that *R. gorgonias* is an AM-competent species, consistent with its nearly complete repertoire of symbiosis genes.

**Figure 2.**
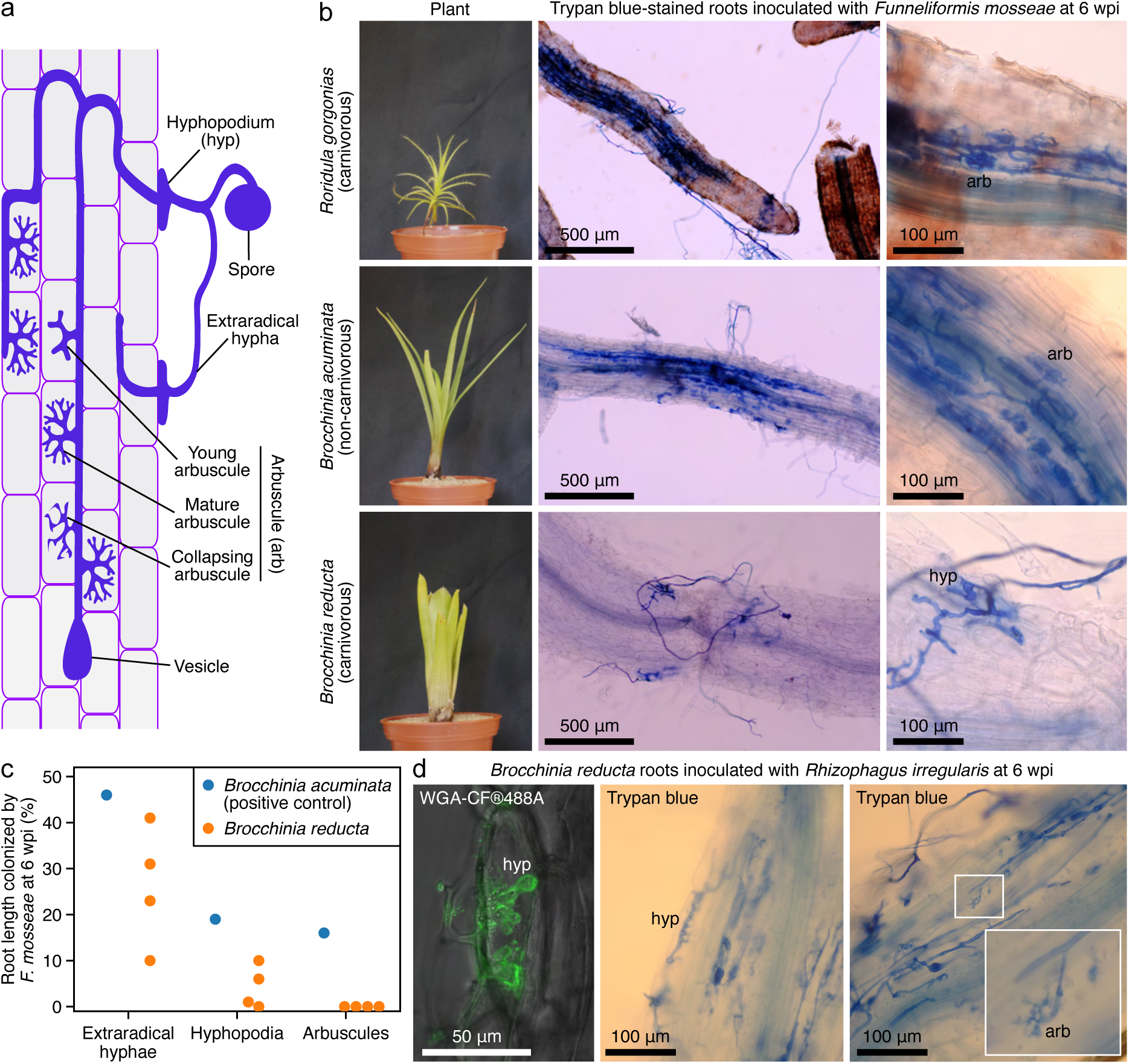
AM inoculation assays in *Roridula* and *Brocchinia*. **a**, Schematic depicting AM colonization and typical AM fungal structures. **b**, Representative images of plants and roots showing *Funneliformis mosseae* colonization. **c**, Quantification of AM symbiotic structures. **d**, Representative microscopy images of *Rhizophagus irregularis* colonizing *Brocchinia reducta* roots. Supplementary Fig. 3 displays all additional arbuscules encountered in *B. reducta* roots. wpi, weeks post inoculation.

While *Roridula* comprises exclusively carnivorous species, *Brocchinia* includes both carnivorous and non-carnivorous species. This diversity enables the comparison of AM status between closely related species with and without plant carnivory. To take advantage of this, we examined the carnivorous *B. reducta* and a non-carnivorous relative *B. acuminata* for AM colonization ability. *Brocchinia acuminata* exhibited well-colonized infection sites harbouring fully developed arbuscules at six wpi. In contrast, several *B. reducta* plants examined showed extraradical hyphae and hyphopodia, but arbuscules were not observed (Fig. 2b-c).

To explore potential differences in colonization ability among fungal species, we performed additional AM inoculation assays in *B. reducta*, employing another AM fungus, *Rhizophagus irregularis*. The observations were similar to those with *F. mosseae*, but detailed inspections on the rare occurrences of hyphopodia revealed projections into the root epidermis, suggesting difficulties in root entry. In addition, we observed arbuscules, albeit infrequent, small and stunted (Fig. 2d). The dozen stunted arbuscules encountered in the root systems of three *B. reducta* plants are shown in Supplementary Fig 3. In summary, AM competence exists within the genus *Brocchinia*, as evidenced by the non-carnivorous *B. acuminata*. However, the carnivorous *B. reducta* displayed aberrant AM colonization, with infrequent and morphologically altered symbiotic structures. This suggests that *B. reducta* can interact with and be colonized by AM fungi but is unable to establish a successful symbiosis.

Given the mutually exclusive trends observed between AM symbiosis and carnivory, we further investigated the AM status of Stylidium, a plant exhibiting characteristics similar to carnivorous species. *Stylidium*, belonging to the Asterales, was previously proposed to be protocarnivorous based on proteinase activity in its sticky glandular hairs (Darnowski et al. 2006). Recent isotope analyses indicated that *Stylidium* species are unlikely to rely on captured organisms for nutrients (Nge and Lambers 2018). The transcriptome of *Stylidium debile* harboured most symbiosis genes (Supplementary Fig. 1). In line with this, inoculation assays with *R. irregularis* showed good colonization levels and the formation of numerous arbuscules, supporting the idea of *S. debile* being a non-carnivorous AM-competent species (Supplementary Fig. 4).

To confirm the non-AM status of carnivorous plants lacking a majority of symbiosis genes, inoculation assays were performed in the carnivorous species *C. follicularis*, which resulted in no observed symbiotic structures in its roots (Supplementary Fig. 5). The retention rates of bioinformatically identified symbiosis genes tightly correlated with experimentally validated AM status, highlighting the effectiveness of using symbiosis gene presence as a reliable predictor of AM competence. Based on this genotype–phenotype association, we infer that other carnivorous lineages with similarly limited symbiosis gene repertoires are also likely to be AM-incompetent.

### Tissue-specific expression of retained AM-associated genes in *Cephalotus follicularis*

Despite its AM incompetence, we detected 15 symbiosis genes in the carnivorous pitcher plant *C. follicularis*. To infer their functions, we examined previously published tissue-specific transcriptome data (Fukushima et al. 2017; Saul et al. 2023). All but two genes, *BLUE COPPER-BINDING PROTEIN1* (*BCP1*) and one *CYTOCHROME P450* (*P450*) gene, were expressed in the analyzed tissues including aerial parts (Fig. 3), suggesting their roles in non-AM-related functions. The two tandemly-duplicated *CHITINASE* orthologs (Cfol_v3_20842 and Cfol_v3_20843) were expressed specifically in the trapping pitcher with the highest expression levels in the upper and lower pitcher walls, where multicellular large glands are located on the inner surface (Juniper et al. 1989). The chitinase enzyme encoded by Cfol_v3_20843 has been confirmed to be secreted into the digestive fluid (Fukushima et al. 2017), suggesting a co-option for digestive physiology to degrade arthropod chitin. In addition, two of the eight *P450* orthologs (Cfol_v3_12140 and Cfol_v3_12147) showed pitcher-preferential expression. They belong to a cytochrome P450 subfamily involved in regulation of disease resistance and apocarotenoid biosynthesis (Wakabayashi et al. 2021; Wang et al. 2022). *DUF538* (Cfol_v3_03686) also displayed higher expression levels in pitchers compared to flat leaves. While it is tempting to suggest that symbiosis genes may have been repurposed for carnivory, these particular genes identified via phylogenomics have not been functionally characterized in AM symbiosis. This highlights the need for future experimental studies to assess their roles in AM symbiosis in non-carnivorous plant species and to explore their potential co-option in carnivorous plants.

**Figure 3.**
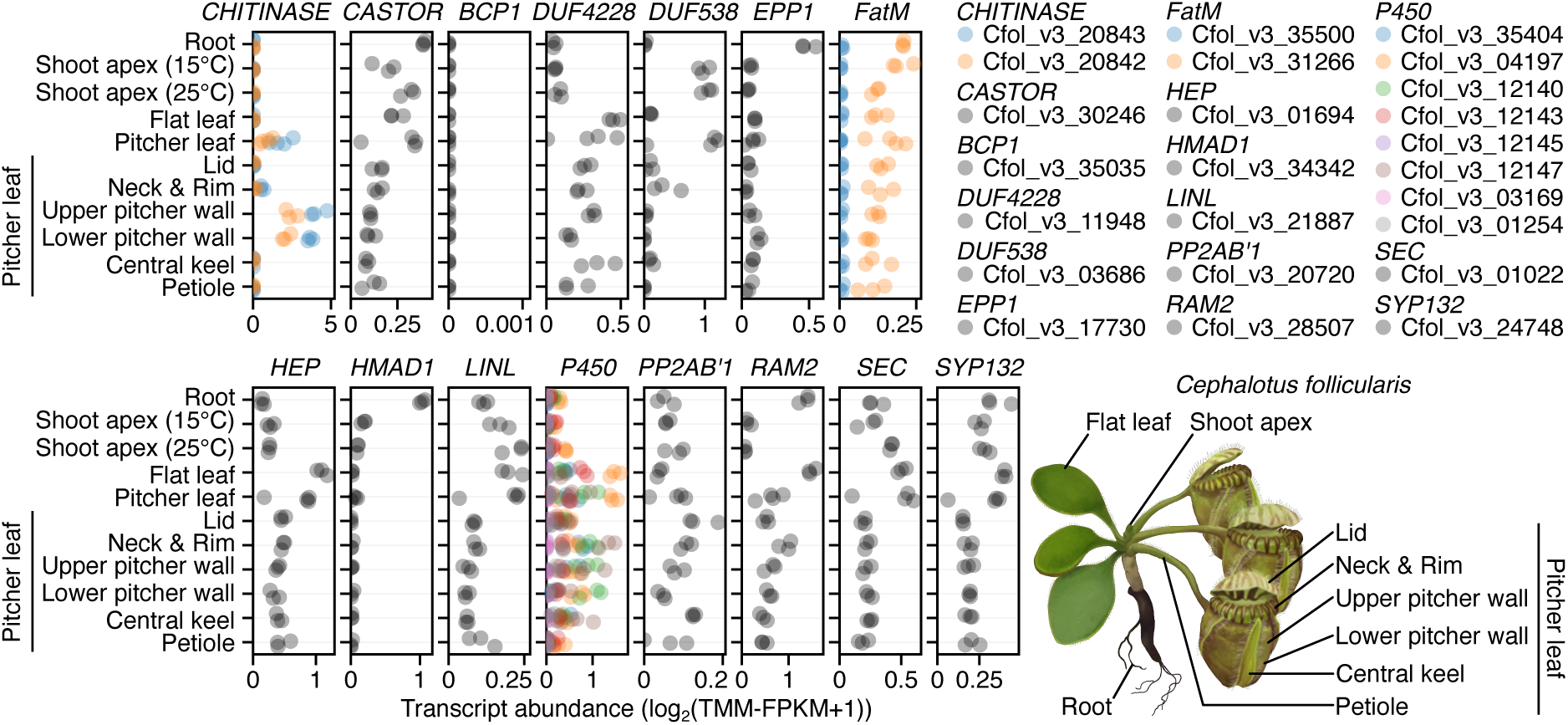
Tissue-specific expression of retained AM-associated genes in the carnivorous pitcher plant *Cephalotus follicularis*. Each point represents a replicate from RNA-seq experiments. Illustration by Shun Anzai.

## Discussion

Our bioinformatic analysis, spanning multiple origins of carnivory and ecological habits, showed that the AM trait has been lost multiple times in carnivorous plant lineages. An earlier phylogenomic study examined only the genome of a single carnivorous plant, *Utricularia gibba* (Bravo et al., 2016). Although *U. gibba* was categorized as non-AM, ambiguity remains regarding the relationship between AM competence and carnivory. This uncertainty arises not only from the lack of data on other carnivorous lineages but also because *U. gibba* is an aquatic herb that lacks roots, a lifestyle incompatible with AM symbiosis. Therefore, our findings establish the mutually exclusive trend between AM competence and carnivory.

Some families in the Caryophyllales such as the Amaranthaceae and Caryophyllaceae are known to be non-AM (Lambers and Teste 2013), and a previous study analyzed two genomes from this order, confirming the absence of symbiosis genes (Bravo et al. 2016). However, we did not anticipate the near-complete absence of symbiosis genes across the entire order of our dataset including 23 families. Some members in the Caryophyllales have been reported to be AM-competent (Maherali et al. 2016), mainly based on information from compilation lists (Wang and Qiu 2006; Akhmetzhanova et al. 2012). However, these compilation lists have been criticized for having misdiagnosed AM-types, which are perniciously recycled in subsequent studies (Brundrett and Tedersoo 2019). Our review of the literature found no convincing evidence of AM colonization and arbuscule formation in the Caryophyllales. Our findings suggest most, if not all, Caryophyllales are non-AM. Conceivably, the early loss of AM symbiosis provided the potential for the development of alternative nutrient acquisition strategies to evolve in this order. This may include the evolution of ectomycorrhizal symbiosis in several Caryophyllales families (Tedersoo and Brundrett 2017), and may also be behind the evolution of carnivory, which reaches its maximum genus diversity in this order (Fleischmann et al. 2018).

In the carnivorous Ericales, symbiosis genes were mostly undetected in the species in the Sarraceniaceae and mostly detected in *Roridula gorgonias*. With experimental evidence of AM colonization, *R. gorgonias* emerged as a remarkable outlier to the non-AM trend observed in carnivorous plants. Mycorrhizal symbiosis in *Roridula*, a genus with two South African species (POWO, 2024), has been largely unexplored, but one century ago, botanist Rudolf Marloth noted in his Flora of South Africa that *Roridula* roots are “surrounded by the loose web of a mycorrhiza” (Marloth 1925). The biological significance of the AM competence in *Roridula* can at present only be speculated based on its unique traits compared to other carnivorous species. Although *R. gorgonias* is almost entirely covered in sticky hairs that efficiently trap insects, it lacks digestive enzymes and relies on symbiosis with a Hemipteran insect that consumes and digests the trapped prey to subsequently pass the digested nutrients as droppings to the plant (Anderson and Midgley 2002). Thus, *Roridula* must share the prey-derived nutrients with symbiotic insects, and this even more unique nutritional mode among carnivorous plants may have led to the maintenance of AM symbiosis as a supplemental source of nutrients.

A total of 50 symbiosis genes were detected in the transcriptome of the carnivorous bromeliad *B. reducta*. Mycorrhizal symbiosis in the genus *Brocchinia*, which comprises 20 species from South America (POWO, 2024), remains largely unknown. Only *B. reducta* corresponds to a confirmed carnivorous species (Givnish et al. 1984). Its closest relative within the genus is *B. hechtioides* (Givnish et al. 1997), but carnivory in this species has not been thoroughly studied. The family they belong to, the Bromeliaceae, is highly diversified and, the sequencing of internal transcribed spacer 1 (ITS1) revealed its species can host diverse mycorrhizal fungal assemblages in their roots (Leroy et al. 2021). One of the most known terrestrial bromeliads, pineapple (*Ananas comosus*), is capable of hosting AM fungi and form arbuscules (Moreira et al. 2015). In our inoculation assays, we observed fully developed arbuscules in roots of the non-carnivorous *B. acuminata*, demonstrating that AM symbiosis does occur in the genus. However, while roots of the carnivorous *B. reducta* were able to interact with AM fungi by forming occasional hyphopodia, intraradical hyphae, and arbuscules, symbiotic structures were malformed. The malformed hyphopodia resembled those described in mutants of symbiosis genes such as *VAPYRIN* in other plant species (Reddy D. M. R. et al. 2007; Pumplin et al. 2010). The few arbuscules encountered were all small and stunted, resembling collapsing arbuscules. Although collapsing arbuscules are common in AM-competent species, they usually coexist with young and fully developed arbuscules. Environmental factors such as nutrient availability, substrate and temperature can quantitatively affect colonization patterns, but the inability of *B. reducta* to form fully developed arbuscules suggests a potential genetic impairment. Thus, *B. reducta* colonization patterns may reflect an incipient loss of symbiosis gene functionality, leading to a diminished ability to establish successful AM symbiosis. The notion of an incipient loss of symbiosis genes is further supported by the mutually exclusive pattern between carnivory and AM symbiosis, as *B. reducta* represents the youngest carnivorous lineage (Fleischmann et al. 2018). The genus *Brocchinia* itself is young, having emerged around 1.9 million years ago, while other carnivorous genera are older, with *Nepenthes*, for example, originating nearly 85 million years ago. These findings invite further genomic studies across *Brocchinia* species engaging in different lifestyles.

Overall, our findings indicate that AM symbiosis and carnivory do not tend to co-occur, possibly due to their overlapping functions. However, the temporal relationship between the loss of symbiotic ability and the acquisition of carnivory does not always appear to be the same. For instance, in lineages like *Brocchinia*, AM symbiosis appears to have been lost upon the evolution of carnivory. By contrast, in the Caryophyllales, AM symbiosis was lost first, and carnivory evolved later. This illustrates that diverse evolutionary paths can lead to the convergence of similar trait combinations.

Our analysis also showed non-random patterns of symbiosis gene retention. In the Poales, symbiosis genes were mostly absent from the sedges *Carex littledalei* and *Rhynchospora tenuis* (Cyperaceae) and *Juncus effusus* from the related family Juncaceae. Sedges are known to develop dauciform roots specialized for phosphate scavenging which are analogous to the cluster roots of the Proteaceae (Shane et al. 2006). The symbiosis genes retained in these species, such as *ALGINATE LYASE 2* (*ALYASE2*) and *HYPOTHETICAL PROTEIN 7* (*HYP7*) may be pleiotropic, with additional unknown roles beyond symbiosis. In the Ericaceae, we identified a unique set of retained symbiosis genes that differ from those found in the above-mentioned monocots. Species in this family are known to develop ericoid mycorrhizal symbiosis, where non-AM fungi form intracellular hyphal coils in epidermal cells (Smith and Read 2008; Leopold 2016). The symbiosis genes retained in *Rhododendron delavayi* and *Vaccinium darrowii* may regulate both the AM and the ericoid mycorrhizal symbioses while the genes absent in the two Ericaceae species may play specific roles in the AM symbiosis. An earlier survey of some symbiosis genes in the transcriptome of *Rhododendron fortunei* showed *SYMRK*, *CCaMK* and *CYCLOPS* to be present (Radhakrishnan et al. 2020). Our data confirms the presence of these genes in the Ericaceae and expands the known repertoire of symbiosis genes in this family (e.g., *ARK2*, *CASTOR* and *VAPYRIN*). It would be fascinating if retained symbiotic genes were repurposed for alternative nutritional strategies, much like CHITINASE, which is encoded by a symbiotic gene, and is secreted into the digestive fluids of the carnivorous plant *Cephalotus*.

In the Lamiales, symbiosis genes were undetected in the Orobanchaceae parasites *Phtheirospermum japonicum* and *Striga asiatica*. Interestingly, the genes were present in the non-parasitic *Lindenbergia philippensis*, which belongs to the same family. These findings suggest a loss of the AM trait in parasitic plants as well as dauciform-forming and ericoid mycorrhizal plants. However, to confirm a widespread absence of symbiosis genes across these different plant guilds, dedicated analyses of each respective trait are necessary, particularly in light of our discovery that *Roridula* serves as an outlier AM species among typically non-AM carnivorous plants.

AM symbiosis and carnivory are both evolutionarily successful strategies for nutrient acquisition. Understanding the switch between these strategies not only enhances our knowledge of carnivory and AM symbiosis but also elucidates how plants fine-tune nutrient acquisition for maximal fitness.

## Methods

### Species set

We surveyed for coding DNA sequence (CDS) fasta files among publicly available genomes in the target orders Poales, Oxalidales, Caryophyllales, Ericales and Lamiales, totalling 82 genomes. One species per genus was included, except in carnivorous plants, where all available were used. In the case of the Poaceae, which has many genomes available, one genome per subfamily was included. Likewise, we surveyed and collected transcriptomes in all families harbouring carnivorous plant species and also in species belonging to families not represented in the genome dataset within the five target orders, totalling 90 transcriptomes. We supplemented this with newly generated genomes and transcriptomes from a total of 15 species, many of them being carnivorous. These data were generated as part of a separate study, but transcriptome assemblies and genome-based coding sequence sets needed to reproduce this study are fully available in Supplementary Data. Additionally, we included 40 genomes outside of the focal orders, aiming for the representation of all angiosperm orders with genomes available. This resulted in a final dataset of 123 genomes and 103 transcriptomes. File sources and benchmarking universal single-copy orthologues (BUSCO) scores are listed in Supplementary Table 2.

### Transcriptome assembly and contamination removal

Publicly available RNA-seq data were retrieved and preprocessed using AMALGKIT v0.9.34 (https://github.com/kfuku52/amalgkit), which internally uses parallel-fastq-dump v0.6.7 (https://github.com/rvalieris/parallel-fastq-dump) and fastp v0.23.2 (https://github.com/OpenGene/fastp). Transcriptome assembly was performed using up to 30 Gb of RNA-seq reads per species with Trinity v2.13.2 (https://github.com/trinityrnaseq/trinityrnaseq). Open reading frames (ORFs) were identified using TransDecoder v5.5.0, with a minimum length threshold of 50 bp (-m 50) (https://github.com/TransDecoder/TransDecoder). The longest ORFs among isoforms were selected using the ’aggregate’ function in CDSKIT v0.10.9 (https://github.com/kfuku52/cdskit). To prevent false detection of potentially contaminated sequences, taxonomic assignment was conducted using MMseqs2 v13.45111 (https://github.com/soedinglab/MMseqs2), and sequences assigned to inconsistent phyla were removed. Finally, the remaining sequences were compiled to generate representative CDSs. Translated CDSs were analyzed using RPS-BLAST v2.13.0 against Pfam-A families (released on November 25, 2020) with an E-value cutoff of 0.01 to determine protein domain architectures.

### Species tree inference

The species tree was inferred as described previously (Saul et al. 2023), utilizing a total of 1,614 single-copy genes identified by BUSCO v5.3.2 (https://gitlab.com/ezlab/busco) with the Embryophyta dataset in OrthoDB v10 (embryophyta_odb10). In-frame codon alignments were generated by first aligning translated protein sequences using MAFFT v7.520, followed by trimming and back-translation with TrimAl v1.4.1 (https://github.com/inab/trimal) to obtain in-frame codon alignments. For each single-copy gene, nucleotide and protein maximum-likelihood (ML) trees were constructed using IQ-TREE v2.2.2.7 (https://github.com/iqtree/iqtree2) with the GTR+R4 and LG+R4 models, respectively. The collection of 1,614 single-copy gene trees was then subjected to coalescence-based species tree inference using ASTRAL v5.7.3 (https://github.com/smirarab/ASTRAL). Additionally, concatenated alignments were used as input for nucleotide- and protein-based ML tree inference with the aforementioned substitution models. *Amborella trichopoda* was used as the outgroup for rooting. The protein-based ML tree was employed as the species tree for subsequent phylogenetic reconciliation analyses.

### Symbiosis gene set

Symbiosis genes used here were largely identified in a previous phylogenomics study (Bravo et al. 2016). Some genes were included as they were shown to have an AM-associated pattern in later studies, including *CCaMK, LIN/LINL* (Radhakrishnan et al. 2020), and *LysMe* (Yu et al. 2023). In total, this corresponds to 75 genes (Supplementary Table 3). Symbiosis genes were identified in our dataset by searching for the orthogroups containing the *Medicago truncatula* orthologs, whose IDs are provided in (Bravo et al. 2016). Gene IDs of other model species were either obtained from the relevant literature when the genes were characterized or, when not characterized, from the Ortholog Search function in the Tomato Symbiotic Transcriptome Database (https://efg.nju.edu.cn/TSTD/, Zeng et al., 2023).

### Orthogroup tree inference

To ensure robust conclusions, we used two different approaches for orthogroup classification: a phylogenetic approach and a graph-based approach. For phylogenetic orthogroup extraction, a TBLASTX search (v2.14.0, (Camacho et al. 2009)) was first performed against the 229 CDS sets using preselected symbiosis genes as queries, with an E-value cutoff of 0.01 and a minimum BLAST hit coverage of 25% on the query sequence, to obtain up to 5,000 homologous sequences. Next, an ML phylogenetic tree was inferred using IQ-TREE with the GTR+G4 nucleotide substitution model, following the generation of in-frame codon sequence alignments as described above. Trees containing more than 2,000 sequences were inferred with the ‘--fast’ option. After species-tree-guided gene tree rooting (Fukushima and Pollock 2020), which internally utilizes NOTUNG v2.9 (https://www.cs.cmu.edu/~durand/Notung/), the phylogenetic orthogroups containing the BLAST query sequences were finally extracted using NWKIT v0.11.10 (https://github.com/kfuku52/nwkit) with the ‘--orthogroup’ option. The graph-based orthogroup inference was conducted using SonicParanoid v2.0.5a2 (https://gitlab.com/salvo981/sonicparanoid2) with representative CDS sets from 229 species, resulting in a total of 110,791 orthogroups from which symbiosis gene orthogroups were identified. ML trees were inferred using the same procedure. All orthogroup trees obtained from the two methods were subjected to the evaluation of tree topology confidence using ultrafast bootstrapping (--ufboot 1000) and optimized through hill-climbing nearest neighbor interchange (-bnni) in IQ-TREE. The ML tree was then used as the starting gene tree for GeneRax v2.0.4 (https://github.com/BenoitMorel/GeneRax) to generate a rooted, species-tree-aware orthogroup tree.

### Symbiosis gene occurrence

Using the SonicParanoid and BLAST-based trees, the occurrence of the different symbiosis genes was assessed in the species within the dataset. In the case of SonicParanoid trees, different symbiosis genes were occasionally encountered in a common orthogroup. The situation where one symbiosis gene was split into two orthogroups also occurred. BLAST-based trees were also commonly composed of the target symbiosis gene and their close non-symbiosis paralogues. In these cases, symbiosis genes were discerned by searching the known *M. truncatula* and rice symbiosis gene orthologs. The subtrees containing them, having an *A. trichopoda* as the basalmost ortholog, were individualized and all the genes contained in these subtrees were considered part of the symbiosis gene group, irrespective of lineage-specific duplications, genes coding for seemingly incomplete proteins or genes with misplaced phylogenetic positions. Symbiosis genes were considered as present if they were present in either the SonicParanoid tree or their BLAST-based tree counterparts. Trees can be found in Supplementary Data.

### Genome-wide analysis of convergent gene losses

To study genes commonly lost in carnivorous plants, we determined orthogroups that were commonly absent in all the carnivorous plant genomes and transcriptomes in the Oxalidales and Lamiales, specifically, *C. follicularis* (Oxalidales), and all species in the genera *Byblis*, *Genlisea, Pinguicula* and *Utricularia* (Lamiales); and present in the non-carnivorous genomes of *Averrhoa carambola* (Oxalidales), *Andrographis paniculata* and *Erythranthe guttata* (Lamiales).

### AM inoculation assays

Young *Brocchinia reducta*, *Brocchinia acuminata*, *Roridula gorgonias* and *Stylidium debile* plants were obtained from a commercial carnivorous plant nursery (Gartenbau Thomas Carow, Nüdlingen, Germany). *Cephalotus follicularis* plants were maintained in tissue cultures (Fukushima et al. 2017). At least three plants of a given species were analysed per experiment with the exception of *B. acuminata*. In this case, we procured a single individual. Roots were carefully washed before transferring plants to pots with autoclaved sand as substrate. AM fungal inoculum was applied. 1,000 spores per pot of the AM fungal species *R. irregularis* (Agronutrition, France) were used as inoculum. In the case of *F. mosseae*, 1 ml of crude inoculum (MycAgro Lab, France) was applied. Plants were grown in the greenhouse for six weeks under a low phosphate fertilization regime administered once a week in the form of 10 μM KH_2_PO_4_.

### AM fungal staining and microscopy

At six wpi, roots were harvested and stained with Trypan blue, largely taking place as described previously (Montero et al. 2021), with an exception for *R. gorgonias* dark roots, where clearing step consisted in 20% (w/v) KOH replaced every other day in a span of a week. Roots were mounted in glass microscope slides, observed in an Olympus CX21 microscope and images were acquired using an affixed FUJIFILM X-T2 camera. For confocal microscopy, roots were stained with Wheat Germ Agglutinin (WGA), CF®488A Conjugate (Biotum). Root sections were excised and incubated in 50% (v/v) ethanol for one hour followed by 20% (w/v) KOH for two days, and 0.1 M HCl for two hours. A 0.2 μg mL^-1^ WGA-CF®488A solution in 1× phosphate-buffered saline (PBS, pH 7.4) was added and samples were incubated at 4°C in the dark for one week. Imaging was carried out in a confocal laser scanning microscope (Leica TCS SP5 II, Leica Microsystems, Wetzlar, Germany). WGA-CF®488A was detected with an excitation wavelength of 488 nm, and emitted wavelengths were collected at 492–533 nm. Roots were observed using a 40 × water immersion objective. Image processing took place using Fiji/ImageJ (Schindelin et al. 2012).

## Data availability

All data are available in the article, its supplementary materials, or on Figshare (https://doi.org/10.6084/m9.figshare.23553813).

## Acknowledgments

We thank Christian Fröschel for providing *R. irregularis* inoculum and Joachim Rothenhöfer for assistance in the plant growth facilities. We thank the collaborative team of the Ancistrocladus and Triphyophyllum genome project for prepublication access to the coding sequences of *Ancistrocladus abbreviatus* and *Triphyophyllum peltatum*. Computations were partially performed on the National Institute of Genetics supercomputer. We acknowledge the following sources for funding: the Sofja Kovalevskaja programme of the Alexander von Humboldt Foundation (to K.F.), a Human Frontier Science Program Young Investigators grant RGY0082/2021 (to K.F.), and JSPS KAKENHI 23K20050 (to K.F.).

## Author Contributions

H.M. and K.F. conceived the study. H.M. and M.F. performed the experiments. H.M., M.F. and K.F. analysed the data. H.M. and K.F. wrote the manuscript.

## Competing Interests

The authors declare no competing interests.

## Supplementary Information

### Supplementary Tables

**Supplementary Table 1.** List of 85 orthogroups absent in carnivorous species and present in non-carnivorous species in the Oxalidales and Lamiales.

**Supplementary Table 2.** Sources and BUSCO scores of genomes and transcriptomes employed in our analysis.

**Supplementary Table 3.** List of symbiosis genes, their associated orthogroup IDs, and gene IDs of the representative species *Medicago truncatula* and *Oryza sativa*.

### Supplementary Figures

**Supplementary Figure 1.**
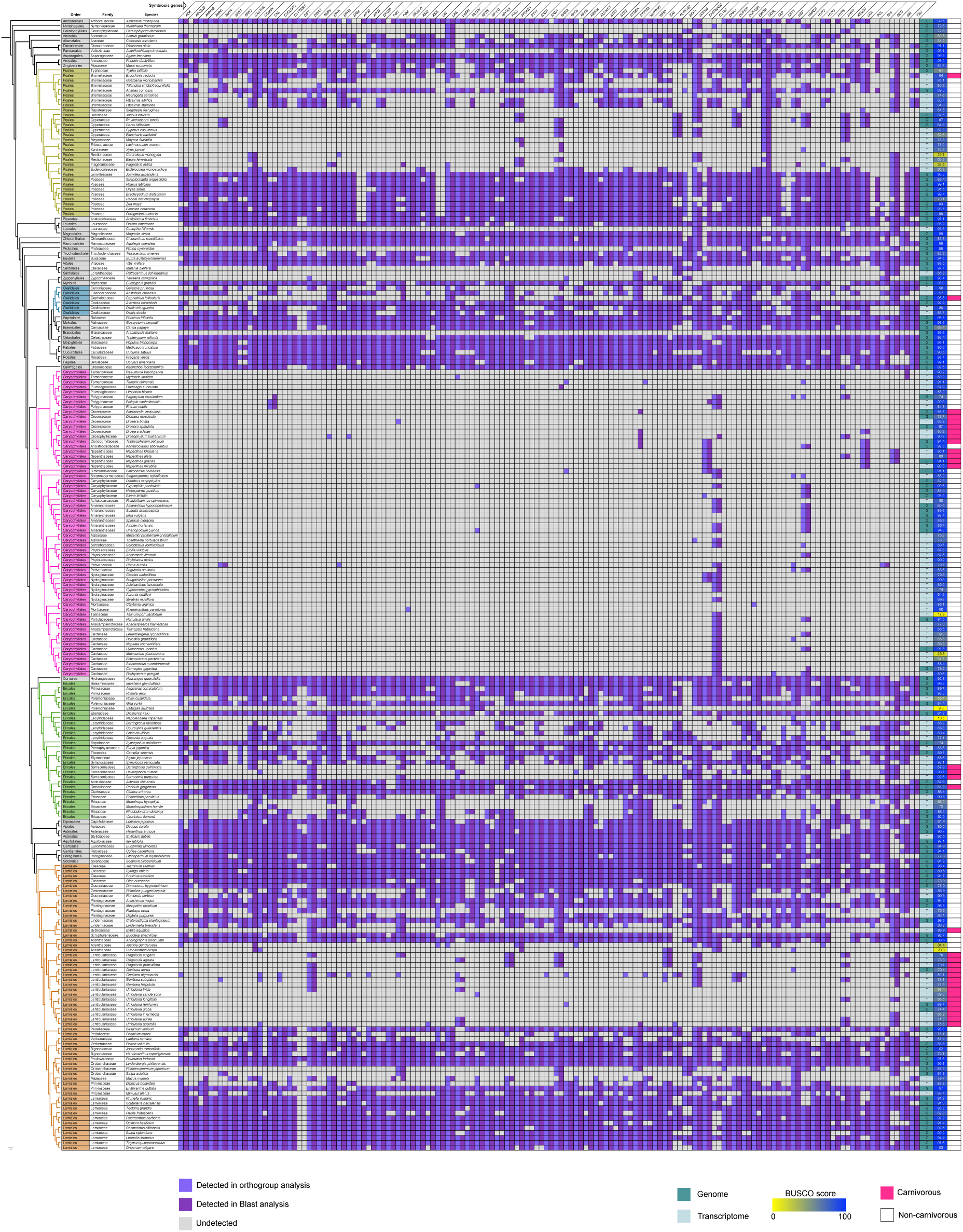
Detail of Figure 1, displaying all species analysed in this study.

**Supplementary Figure 2.**
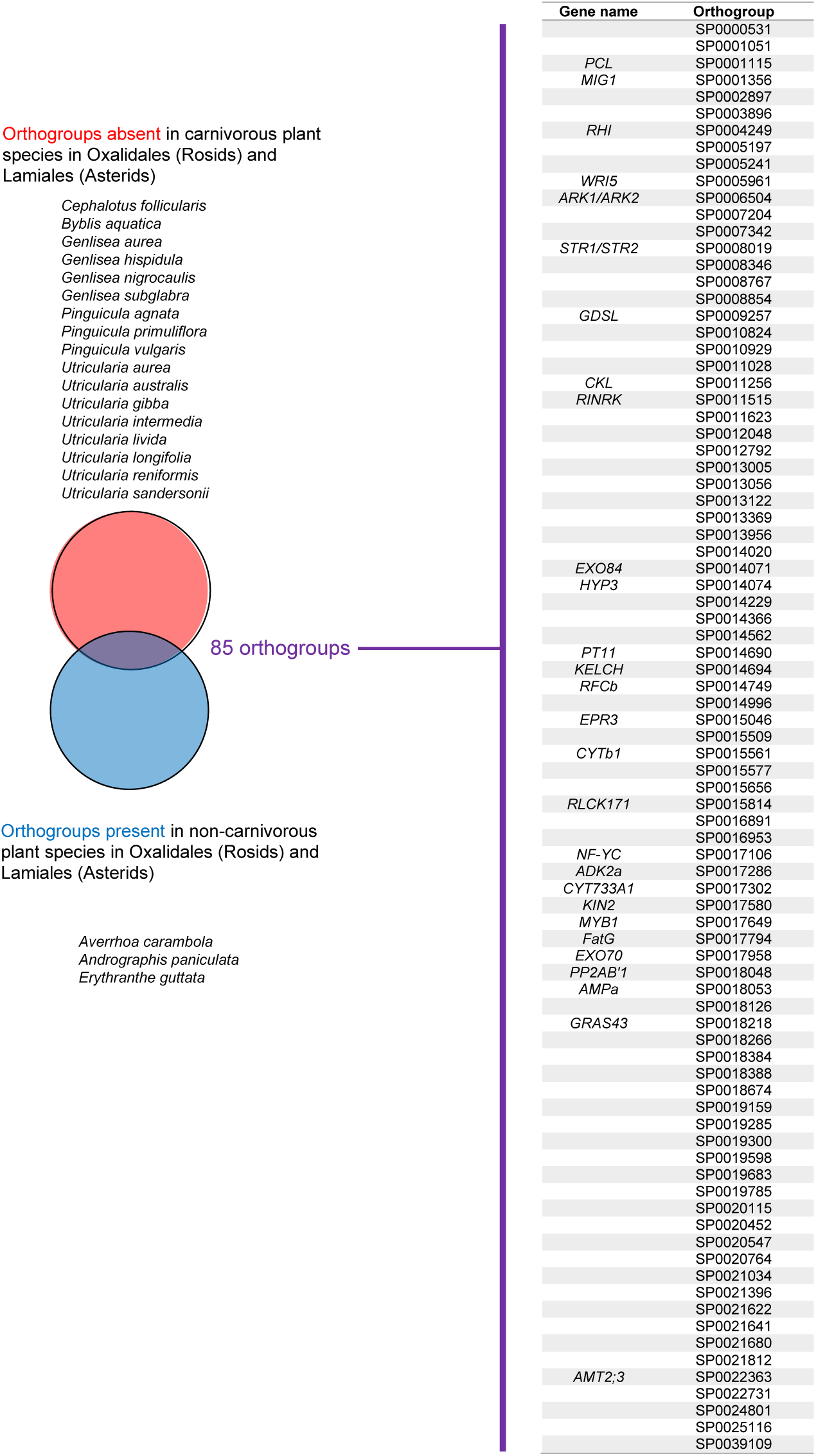
85 orthogroups absent in the genomes and transcriptomes of the carnivorous plant species Cephalotus follicularis, Byblis aquatica, Genlisea aurea, G. hispidula, G. nigrocaulis, G. subglabra, Pinguicula agnata, P. primuliflora, P. vulgaris, Utricularia aurea, U. australis, U. gibba, U. intermedia, U. livida, U. longifolia, U. reniformis and U. sandersonii; and present in the non-carnivorous genomes of Averrhoa carambola, Andrographis paniculata and Erythranthe guttata. Of these, 28 orthogroups correspond to AM-related genes (named next to the orthogroup IDs). Table displaying gene IDs per each species can be found in Supplementary Table 2.

**Supplementary Figure 3.**
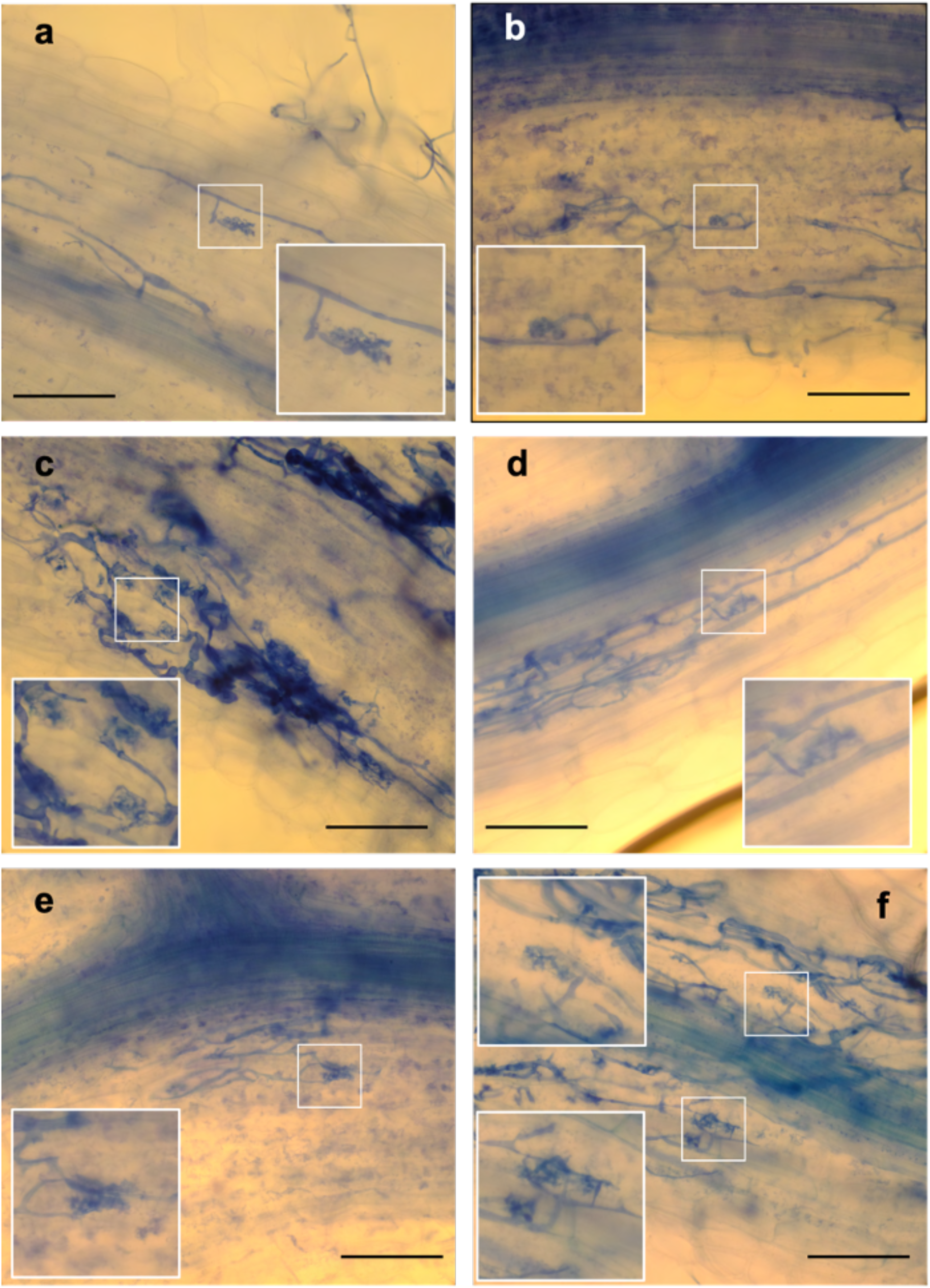
*B. reducta* roots inoculated with *R. irregularis* at six wpi stained with Trypan blue. All root areas colonized and forming arbuscules along the root system of three plants scored are shown. Arbuscules are boxed and images enlarged in insets. a, b, d and e show single stunted arbuscules in sparsely colonized root zones. c and f show multiple stunted arbuscules in root zones with more intraradical hyphae. Scale bar, 100µm.

**Supplementary Figure 4.**
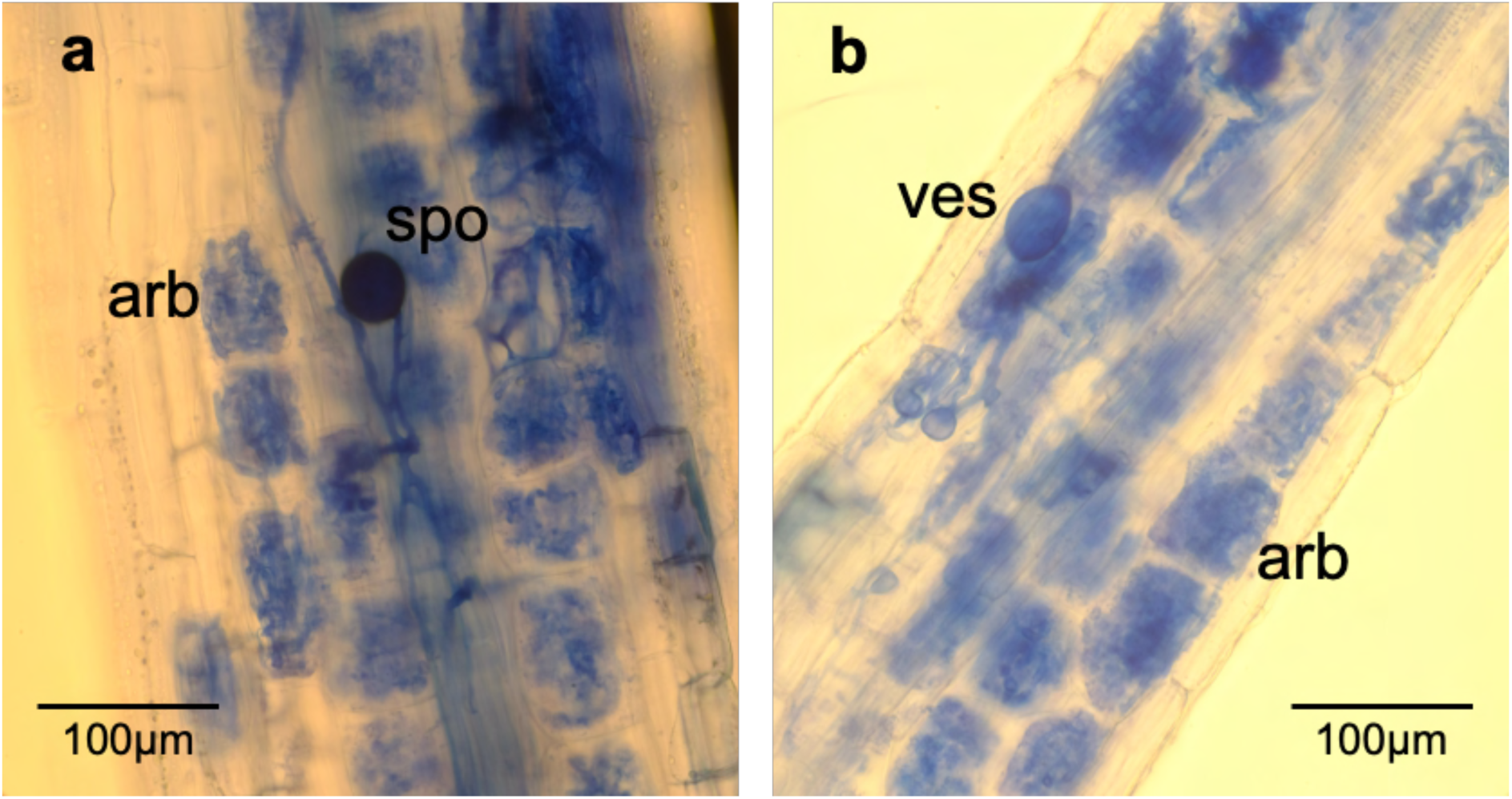
*Stylidium debile* roots inoculated with *R. irregularis* at six wpi stained with Trypan blue. a, arbuscules (arb) and spore (spo). b, arbuscules and vesicle (ves).

**Supplementary Figure 5.**
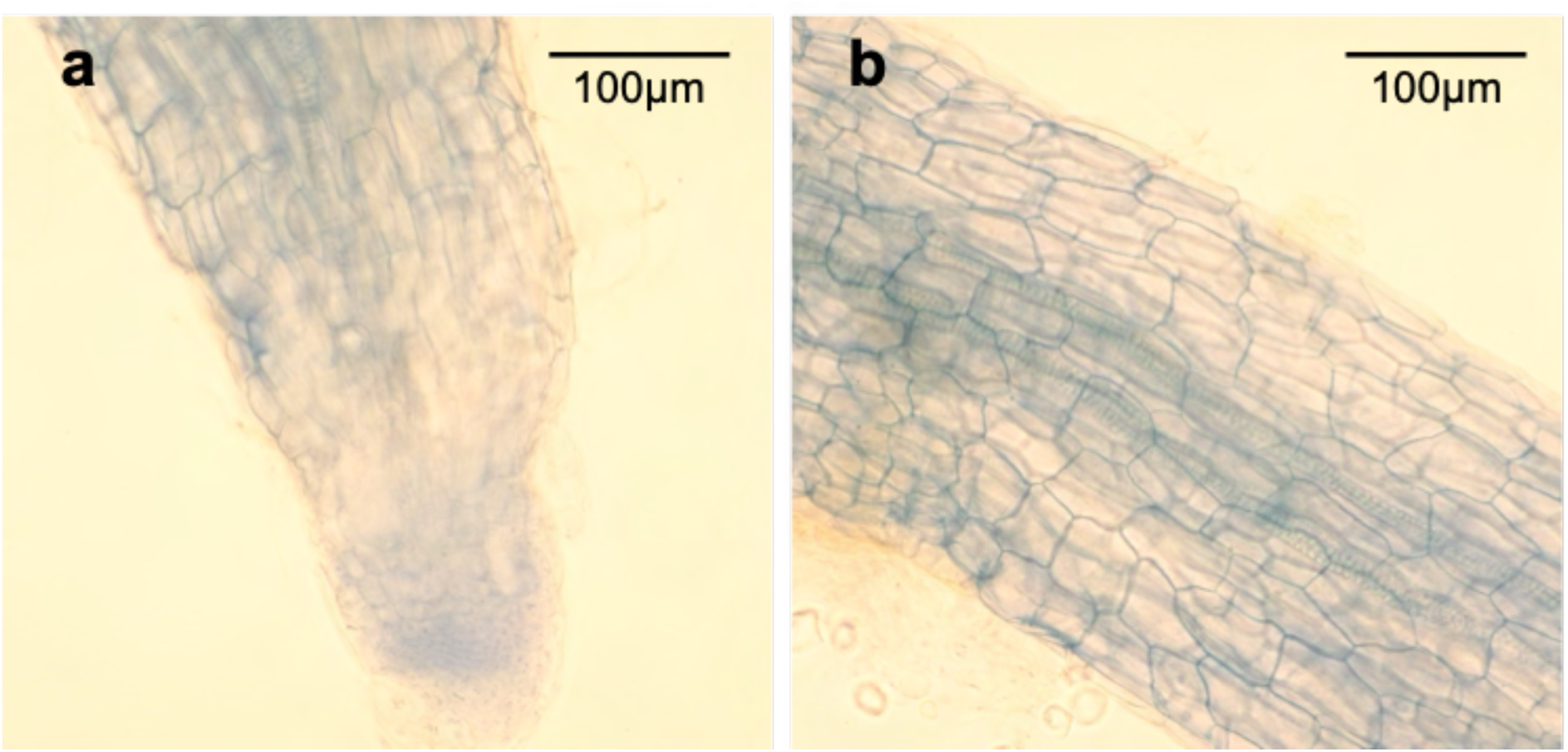
*Cephalotus follicularis* roots inoculated with *R. irregularis* at six wpi stained with Trypan blue. a, root tip. b, root area distal from tip.

## Notes

https://doi.org/10.6084/m9.figshare.23553813

### Competing Interest Statement

The authors have declared no competing interest.

